# Stick Stippling for Joint 3D Visualization of Diffusion MRI Fiber Orientations and Density

**DOI:** 10.1101/2020.06.15.153098

**Authors:** Ryan P. Cabeen, David H. Laidlaw, Arthur W. Toga

## Abstract

This paper investigates a stick stippling approach for glyph-based visualization of complex neural fiber architecture derived from diffusion magnetic resonance imaging. The presence of subvoxel crossing fibers in the brain has prompted the development of advanced modeling techniques; however, there remains a need for improved visualization techniques to more clearly convey their rich structure. While tractography can illustrate large scale anatomy, visualization of diffusion models can provide a more complete picture of local anatomy without the known limitations of tracking. We identify challenges and evaluate techniques for visualizing multi-fiber models and identified techniques that improve on existing methods. We conducted experiments to compare these representations and evaluated them with *in vivo* diffusion MR datasets that vary in voxel resolution and anisotropy. We found that stick rendering as 3D tubes increased legibility of fiber orientation and that encoding fiber density by tube radius reduced clutter and reduced dependence on viewing orientation. Furthermore, we identified techniques to reduce the negative perceptual effects of voxel gridding through a jittering and re-sampling approach to produce a stippling effect. Looking forward, this approach provides a new way to explore diffusion MRI datasets that may aid in the visual analysis of white matter fiber architecture and microstructure. Our software implementation is available in the Quantitative Imaging Toolkit (QIT).

## 1 Introduction

Diffusion magnetic resonance imaging (dMRI) provides a unique probe of water molecule diffusion that can reveal the complex architecture of neural tissue, including crossing, kissing, and bending fibers [1]. This complexity poses a challenge when visualizing dMRI datasets due to the dense and overlapping nature of fiber bundles in the brain [2]. While tractography provides a powerful tool for understanding large scale neural structures, there is high variability across tracking methods and false positives are difficult to avoid [3] and there are limitations to its anatomical accuracy [4]. To help provide a more complete picture of the underlying anatomy and to understand why tracking algorithms fail, it can be useful to instead visualize the underlying voxel models representing local diffusion properties in each voxel [5] [6] [7].

Glyph rendering is a common technique for visualizing local features of dMRI data [8]; in this process, markers or symbols are used to visually encode the diffusion or microstructure properties at various positions of a volume. Previous work has included a variety of useful glyphs [2], including ellipsoids [5] and superquadratics representing tensors [9] [10]. Glyphs have also been used to represent more complex models, such as orientation distribution functions (ODFs) [11], and spherical harmonic representations of fiber orientation distributions (FODs) [12] [13]. Other work has focused on directly visualizing fiber organization through glyph packing [14], and two-dimensional techniques such as line rendering [15] [16] and line stippling [17] [18]. Nevertheless, the visualization of complex fiber configurations remains a major challenge, due to cluttering in high resolution data, dependence on viewing orientation, and the introduction of perceptual artifacts due to the gridding of voxels.

Our primary contribution is the development of a visualization technique that combines several of these existing approaches to better depict crossing fiber configurations with fixels, a term we adopt here to refer to the combination of fiber orientations and associated fiber density or volume fraction (Fig 1) [19]. We propose methods that build on related superquadratic and fiber stippling work by using data-modulated 3D stick glyphs and a stippling effect to more clearly depict crossing fiber orientation and density. We compare these and other multi-fiber glyphs and show how they are affected by image gridding and viewing orientation. We then evaluate this technique by creating visualizations of *in vivo* neuroimaging data with varying voxel resolution and anisotropy to show the benefits of this approach for typical data exploration sessions.

**Fig. 1:**
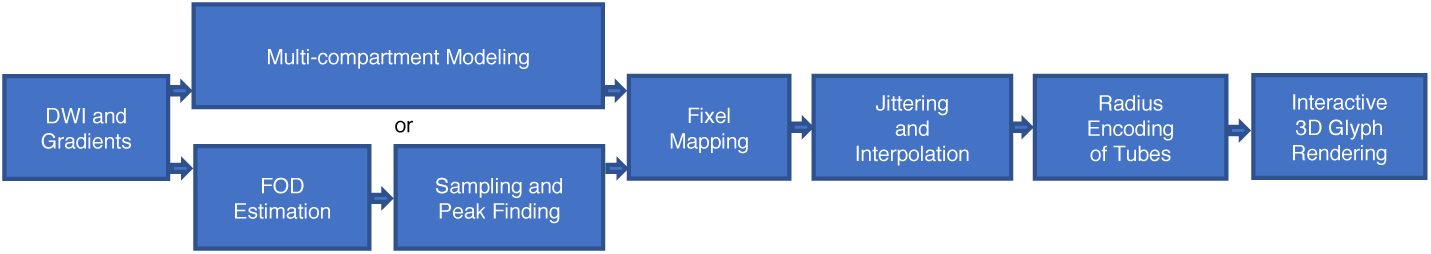
The proposed computational pipeline for creating stick stippling visualizations from diffusion MRI data. The pipeline applies to both multi-compartment models, such as the ball-and-sticks model, as well as FODs represented by spherical harmonics.

## 2 Methods

### 2.1 Diffusion Modeling and the Fixel Representation

We focus here on the visualization of multi-fiber models using a fixel representation that can summarize either FODs [20] or multi-compartment models such as the ball-and-sticks model [21]. Each fixel consists of a fiber orientation and its associated fiber density, which are extracted as follows. For FOD modeling, fiber orientations are extracted by sampling the distribution on a four-fold subdivision of an spherical icosahedral mesh. From this, fiber orientations are extracted from the local maxima on the mesh, and fiber density is taken by the peak density. FOD fixels further processed to remove duplicates at antipodal points and local ridges using hierarchical clustering with a sine angle distance. For multi-compartment modeling, fixels are directly derived from the principal direction and volume fraction of each compartment. While previous work has used fixels primarily for FODs [22], we more generally use fixel density to represent either volume fraction or density, depending on the underlying model.

More formally, the fixels of the *i*-th voxel are given by 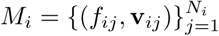, with fixel count *N*, density *f*, and orientation *v*. Our approach also requires interpolation for jittering fixels in a neighborhood around each voxel, and for this, we use a kernel regression framework [23] [24], which estimates the fixels 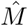 at an arbitrary position *p* from a neighborhood of voxels 𝒩:

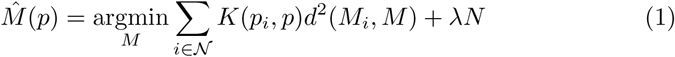

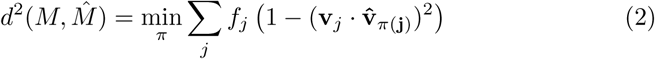

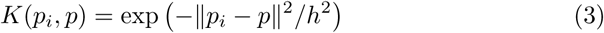

given spatial bandwidth *h*, and regularization *λ* = 0.99. While previous work has used this with ball-and-sticks modeling, we more generally use this here with fixels obtained from FODs [22] as well. The primary goal here is then to develop improved glyph-based visualization techniques based on this approach, which we describe next.

### 2.2 Fixel Glyph Visualization

The design of glyph visualizations generally includes a number of technical challenges related to visual perception [25] [26] [27] [28], and we identified three major issues specifically related to fixels (Fig. 2). First, the complexity of multi-fiber models often creates cluttered scenes, making it difficult to distinguish important fibers from unimportant ones. Second, the viewing orientation of the scene can reduce the legibility of fixel orientation and density, particularly those parallel to the camera’s viewing direction. Third, the discretization of the voxel grid can break gestalt principles of perceptual organization, making some bundles appear more organized than others, even when they only differ in orientation relative to the viewing angle.

**Fig. 2:**
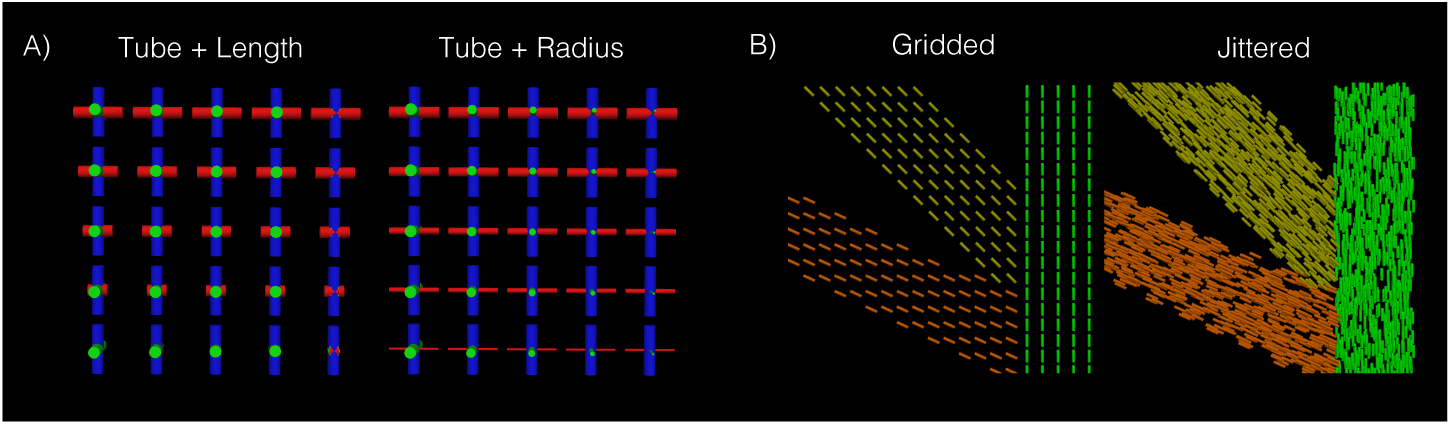
Perceptual challenges for fixel visualization shown in synthetic data. Panel A shows challenges related to viewing orientation, where length and radius encoding of fiber density are compared. Red fiber density decreases from top to bottom, and green fiber density decreases left to right. This illustrates how the density of fibers parallel to the camera view (green) is only legible with radius encoding. Panel B shows challenges related to voxel discretization, where gridded and jittered glyphs are compared. Three bundles were created with equal thickness but with different orientations. This illustrates how gridding can make axis-aligned bundles look more coherent, while jittering can avoid these issues to give an equally coherent appearance to all bundles.

We propose to address these issues by using the 3D shape and layout of fibers to reduce visual ambiguities (Fig. 1). First, we render fixels as tubes (or sticks) to provide a clearer reading of fiber orientation, as shading from the 3D shape can provide depth cues, unlike rendering solid lines. We also color fixels according to the widely used RGB-directional color mapping. Second, we modulate the radius of the tube to reflect the fiber density. The advantage of modulating the radius is that it can be appreciated from any viewing orientation, unlike tube length, which can be illegible when parallel to the viewing direction and can be confused with orientation due to perspective foreshortening. Finally, we render fixels using jittering of 3D glyphs to produce a stippling effect, a process that greatly reduces the effect of gridding by drawing glyphs at uniformly sampled points within each voxel. This approach builds on related work on stippling and superquadratics to visualize fixels. Furthermore. In the following section, we conduct experiments to make comparisons with previously used multi-fiber visualizations and to understand how effectively they address the challenges posed in fixel visualization.

## 3 Experiments and Results

We conducted experiments with three datasets to understand the strengths and limitations of various fixel rendering techniques with regard to how they represent complex anatomy. These datasets were chosen to include voxels of varied resolution and anisotropy in order to show performance across anatomical scales and imaging parameters. In addition to techniques described in the previous section, we compared the results to solid line rendering, tube length encoding, and spherical harmonic FOD rendering when applicable. Our experiments were implemented in the Quantitative Imaging Toolkit (QIT) [29].

### 3.1 Clinical Data Experiment

Our first experiment examined data from a typical clinical acquisition. Diffusion-weighted MR image volumes were acquired from a healthy 34 year old volunteer, conducted on a GE 1.5T scanner with voxel size of 2 mm^3^, matrix size 128×128, and 72 contiguous axial slices. A total of 71 volumes were acquired, with seven T_2_-weighted volumes (b-value 0 s/mm^2^) and 64 diffusion-weighted volumes (b-value 1000 s/mm^2^) and distinct gradient encoding directions. Ball-and-sticks were estimated using MCMC optimization procedure of Behrens et al. [21].

Eight visualizations were generated to depict a coronal slice showing complex interactions among the corpus callosum (CC), and superior longitudinal fasciculus (SLF), cingulum bundle, and corona radiata (Fig. 3). This figure compares gridded and jittered glyph placement, line and 3D tube drawing, and length and radius encodings of fiber density. We found that jittering effectively removed evidence of the underlying voxel grid, and visualization representing fiber density had less clutter. We found that length encoding of volume fraction was only effective in fibers orthogonal to the viewing direction, however, the density of fibers parallel to the view, such as the SLF and cingulum, were only appreciable with radius encoding.

**Fig. 3:**
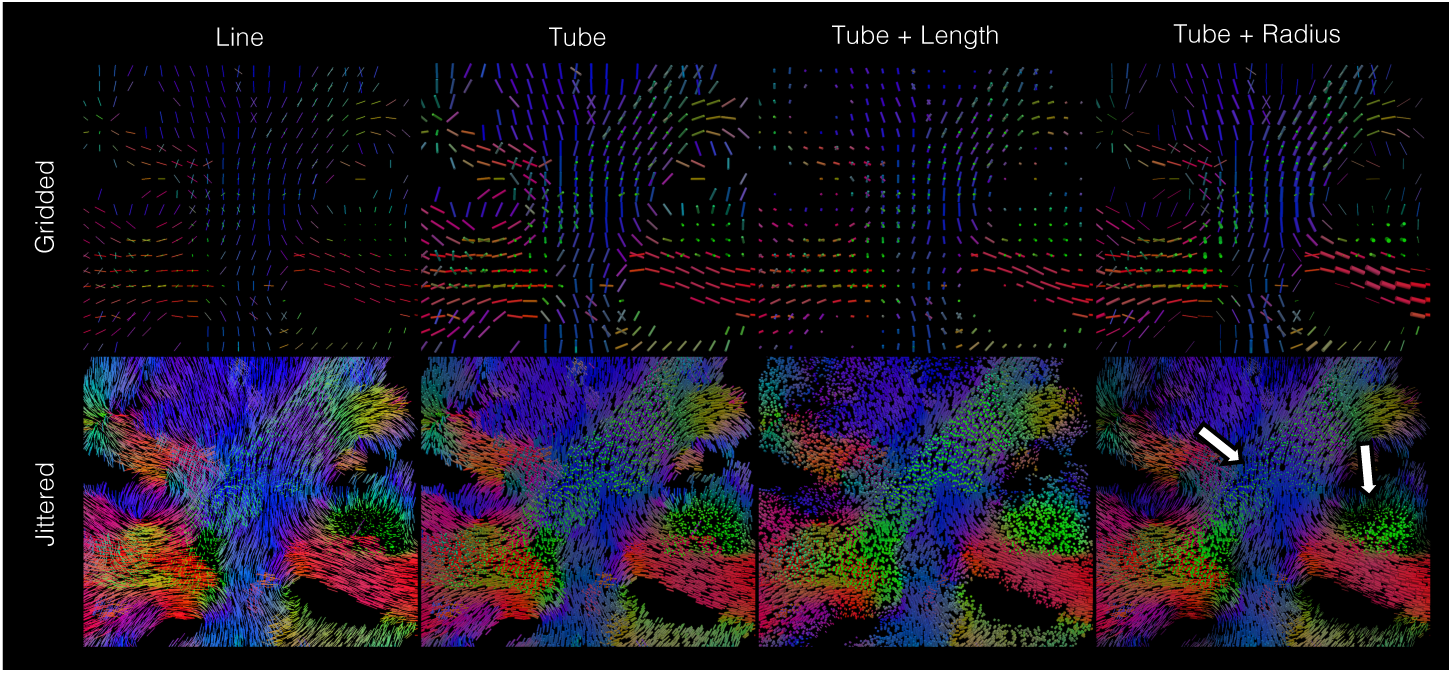
Eight fixel visualizations with clinically feasible data. A comparison of the first and second columns shows how 3D tube rendering more clearly depicts fiber orientation. A comparison of the third and fourth columns show how radius encoding can illustrate variation in fiber density, e.g. the variation in SLF fibers shown in green (white arrows).

### 3.2 HCP Experiment

Our second experiment examined state-of-the-art dataset from the Human Connectome Project ^1^, specifically the single subject dataset with identifier 100307. Diffusion-weighted MR imaging was conducted on a Siemens 3T scanner with voxel size of 1.25 mm^3^, matrix size 145×145, and 174 slices. A total of 288 volumes were acquired, with 18 T_2_-weighted volumes (b-value 0 s/mm^2^) and the remainder distributed among roughly three shells (b-values 1000, 2000, 3000 s/mm^2^) with distinct gradient encoding directions. FODs were estimated using the compartmental modeling approach of Tran et al. [30] using 16 order spherical harmonics. The FODs were discretized using a subdivided icosahedron, and fixels were created from the peak directions and fiber density.

Four visualizations were generated to depict a coronal slice that includes a triple crossing of the corticospinal tract, CC, and SLF (Fig. 4). These visualizations included line rendering, spherical FOD rendering, gridding tubes, and jittered tubes. We found that the high resolution of the data make gridded visualizations more difficult to read, due to the smaller size of fixel glyphs. The tube renderings were more legible than either line or FOD rendering, but jittered tubes provided the greatest legibility. In particular, the jittering more clearly showed triple crossings, density reductions at the white-gray matter interface and the variation in density of the SLF.

**Fig. 4:**
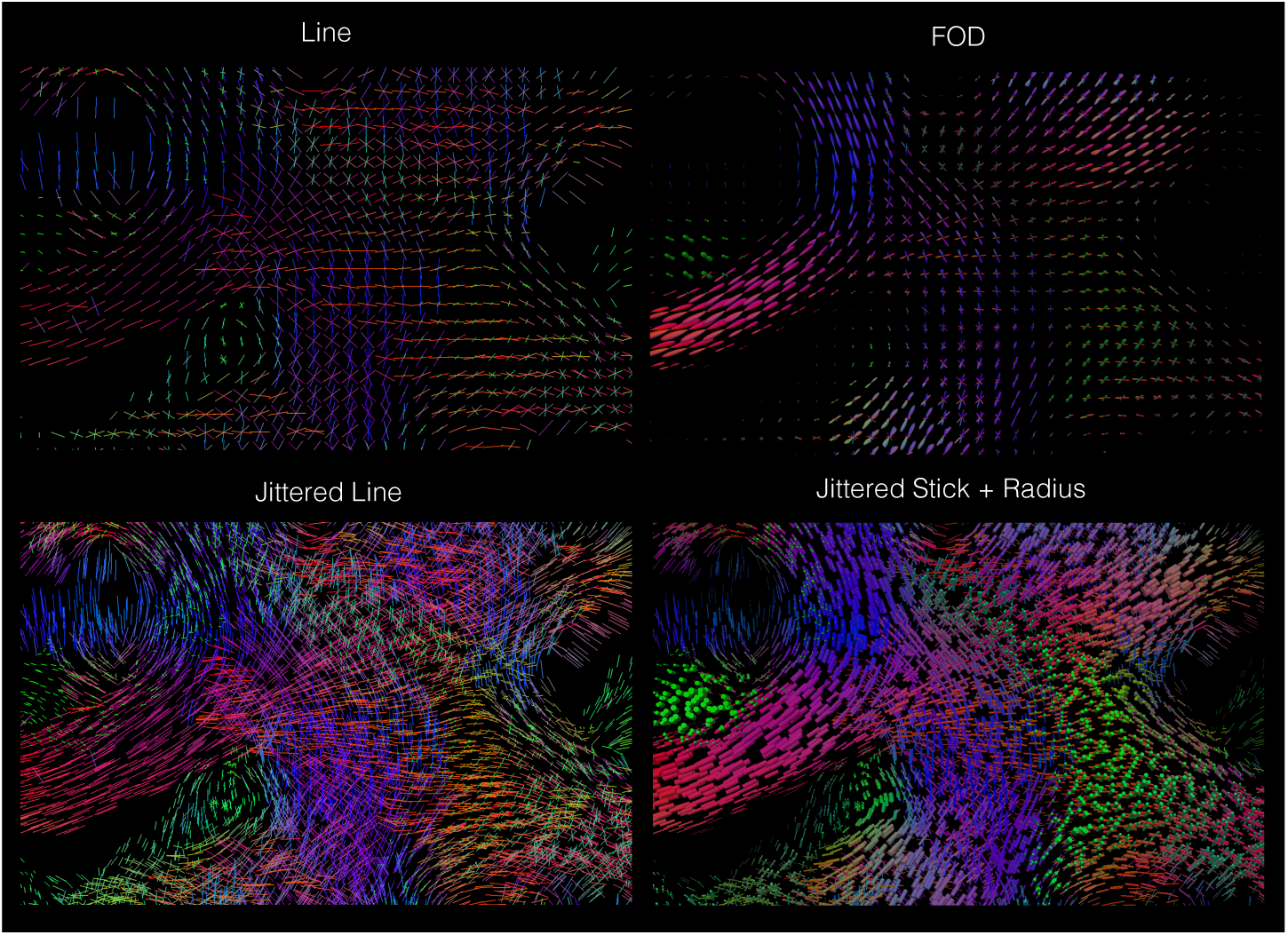
Visualizations of human connectome project data. The top row shows how high spatial resolution can impact visibility in line and ODF rendering. The bottom right shows how jittering can avoid gridding artifacts and improve visibility.

### 3.3 RESOLVE Experiment

Our third experiment examined an advanced presurgical acquisition with highly anisotropic voxels. Diffusion-weighted MR imaging was conducted prior to surgery to treat temporal lobe epilepsy. The RESOLVE DWI sequence was run on a Siemens Prisma 3T scanner with a high resolution coronal in-plane resolution of 0.6mm x 0.6mm with matrix size 272×360. 19 contiguous slices were acquired with thickness 2.1 mm. A total of 56 volumes were acquired, with six T_2_-weighted volumes (b-value 0 s/mm^2^) and 50 diffusion-weighted volumes (b-values 800 and 2000 s/mm^2^) and distinct gradient encoding directions. Ball-and-sticks were estimated using MCMC optimization procedure of Behrens et al. [21] with the continuous exponential model for multi-shell experiments.

Four visualizations were generated to compare fixel visualization of the coronal high in-plane resolution data with an axial slice showing voxel anisotropy (Fig. 5). Each slice was rendered using both gridded and jittered tubes with radius encoding. The coronal slice was found to have fine detail in crossing fibers; however, the axial slice showed perceptual irregularities due to the thick slices.

**Fig. 5:**
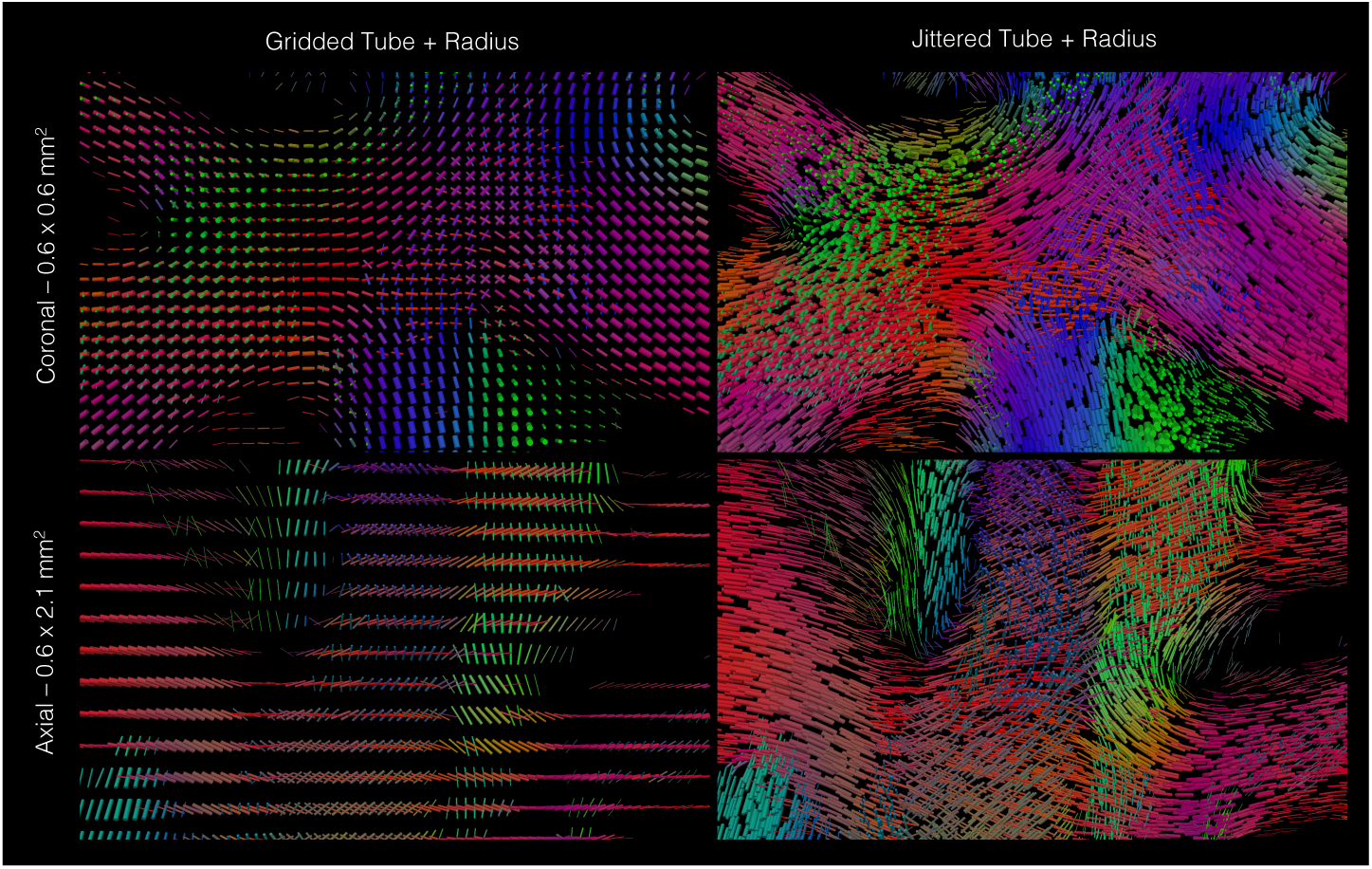
Visualizations of RESOLVE DWI data. The top left shows how high in-plane resolution can greatly reduce the size and visibility of glyphs. The bottom left shows how this introduces large spacing between glyphs when voxels are anisotropic. By contrast, jittered glyphs were found to greatly reduce these negative effects.

Specifically, the anisotropic axial slice showed greater coherence of SLF fibers than CC fibers, due to the greater separation of endpoints of fixels representing the CC. In contrast, the jittered glyphs showed no discrepancy between coronal and axial views, reducing the effects of both voxel gridding and anisotropy.

## 4 Discussion and Conclusions

Our results indicate that when visualizing complex fiber architecture, there are limitations caused by clutter, viewing orientation, and voxel gridding; however, a comparison of glyph visualization techniques showed that these can be mitigated. First, rendering fixels as 3D tube-shaped sticks helped to better indicate orientation through shading and depth cues, and modulating the radius by fiber density helped to reduce clutter and ambiguities based on viewing direction. Second, a stippling effect, produced by glyph jittering and resampling, was found to reduce gridding artifacts, which is perhaps similar to anti-aliasing through randomized glyph placement. Nevertheless, a number of open questions remain. Many applications require the visualization of other per-compartment parameters, e.g. statistical metrics, axon diameter, or myelination, and this requires further work to understand how to effectively encode such parameters with glyphs. There may also be cases where fixels are not a sufficient representation, e.g. when there is fanning or dispersion that may be more apparent by visualizing a complete FOD. There also remain important open problems related to visualizing glyphs in conjunction with tractography [31] and surfaces [32]. Looking forward, the described stick stippling approach may help to guide the design of future imaging studies and aid the development of advanced visualization systems for both human neuroimaging and preclinical imaging [33]. Our software implementation is available online as part of the Quantitative Imaging Toolkit (QIT) ^2 3^.

## Footnotes

^1^http://www.humanconnectome.org

^2^https://cabeen.io/qitwiki

^3^https://resource.loni.usc.edu/resources/downloads/

